# Alcohol consumption is associated with widespread changes in blood DNA methylation: analysis of cross-sectional and longitudinal data

**DOI:** 10.1101/452953

**Authors:** Pierre-Antoine Dugué, Rory Wilson, Benjamin Lehne, Harindra Jayasekara, Xiaochuan Wang, Chol-Hee Jung, JiHoon E Joo, Enes Makalic, Daniel F Schmidt, Laura Baglietto, Gianluca Severi, Christian Gieger, Karl-Heinz Ladwig, Annette Peters, Jaspal S Kooner, Melissa C Southey, Dallas R English, Melanie Waldenberger, John C Chambers, Graham G Giles, Roger L Milne

**Affiliations:** Cancer Epidemiology and Intelligence Division, Cancer Council Victoria, Melbourne, VIC, Australia; Centre for Epidemiology and Biostatistics, Melbourne School of Population and Global Health, The University of Melbourne, Parkville, VIC, Australia; Research Unit of Molecular Epidemiology, Helmholtz Zentrum München, German Research Center for Environmental Health, D-85764 Neuherberg, Germany; Institute of Epidemiology, Helmholtz Zentrum München, German Research Center for Environmental Health, D-85764 Neuherberg, Germany; Department of Epidemiology and Biostatistics, Imperial College London, London W2 1PG, UK; Colorectal Oncogenomics Group, Department of Clinical Pathology, The University of Melbourne, Melbourne, VIC, Australia; Melbourne Bioinformatics, The University of Melbourne, Parkville, VIC, Australia; Department of Clinical and Experimental Medicine, University of Pisa, Pisa, Italy; CESP, INSERM U1018, Univ. Paris-Sud, UVSQ, Université Paris-Saclay, Villejuif, France; Gustave Roussy, Villejuif, France; Klinik und Poliklinik für Psychosomatische Medizin und Psychotherapie des Klinikums Rechts der Isar der TUM, Munich, Germany; German Center for Cardiovascular Research (DZHK), Partner Site Munich Heart Alliance, Munich, Germany; Department of Cardiology, Ealing Hospital, Middlesex UB1 3HW, UK; Imperial College Healthcare NHS Trust, London W12 0HS, UK; MRC-PHE Centre for Environment and Health, Imperial College London, London W2 1PG, UK; National Heart and Lung Institute, Imperial College London, London W12 0NN, UK; Genetic Epidemiology Laboratory, Department of Clinical Pathology, University of Melbourne, VIC, Australia; Precision Medicine, School of Clinical Sciences at Monash Health, Monash University, Clayton, VIC, Australia; Lee Kong Chian School of Medicine, Nanyang Technological University, Singapore 308232, Singapore

**Author notes:** Corresponding author: Pierre-Antoine Dugué, Cancer Epidemiology and Intelligence Division, Cancer Council Victoria.

## Abstract

**Background:** DNA methylation may be one of the mechanisms by which alcohol consumption is associated with the risk of disease. We conducted a large-scale, cross-sectional, genome-wide DNA methylation association study of alcohol consumption and a longitudinal analysis of repeated measurements taken several years apart.

**Methods:** Using the Illumina Infinium HumanMethylation450 BeadChip, DNA methylation measures were determined using baseline peripheral blood samples from 5,606 adult Melbourne Collaborative Cohort Study (MCCS) participants. For a subset of 1,088 of them, these measures were repeated using blood samples collected at follow-up, a median of 11 years later. Associations between alcohol intake and blood DNA methylation were assessed using linear mixed-effects regression models adjusted for batch effects and potential confounders. Independent data from the LOLIPOP (N=4,042) and KORA (N=1,662) cohorts were used to replicate associations discovered in the MCCS.

**Results:** Cross-sectional analyses identified 1,414 CpGs associated with alcohol intake at P<10^-7^, 1,243 of which had not been reported previously. Of these 1,243 novel associations, 1,078 were replicated (P<0.05) using LOLIPOP and KORA data. Using the MCCS data, we also replicated (P<0.05) 403 of 518 associations that had been reported previously. Interaction analyses suggested that associations were stronger for women, non-smokers, and participants genetically predisposed to consume less alcohol. Of the 1,414 CpGs, 530 were differentially methylated (P<0.05) in former compared with current drinkers. Longitudinal associations between the change in alcohol intake and the change in methylation were observed for 513 of the 1,414 cross-sectional associations.

**Conclusion:** Our study indicates that, for middle-aged and older adults, alcohol intake is associated with widespread changes in DNA methylation across the genome. Longitudinal analyses showed that the methylation status of alcohol-associated CpGs may change with changes in alcohol consumption.

## INTRODUCTION

DNA methylation is the addition of methyl groups to the 5’carbon of cytosine in CpG dinucleotides (CpGs) and is thought to play a role in the development of disease through its influence on gene expression and cellular function **(1-4)**. DNA methylation is strongly affected by the underlying genetic DNA sequence **(5)**, sex, age and ethnicity **(5-8)** and is modified by lifestyle factors and environmental exposures such as smoking and adiposity **(9-15)**.

Alcohol consumption is a major lifestyle risk factor contributing to the worldwide burden of disease, responsible for an estimated 2.7 million deaths and 4% of the global burden of disease annually **(16)**. Even modest use of alcohol may increase disease risk, but greatest risks are observed with heavy and long-term drinking. Alcohol consumption is a potentially modifiable risk factor that can be targeted with preventive interventions at both the policy and the individual levels **(17)**. Although there is a plausible relationship between alcohol intake and altered one-carbon metabolism and DNA methylation **(18-20)**, to our knowledge only one large methylome-wide association study (herein referred to as EWAS) of alcohol consumption has been conducted **(21)**. Genes have been reported to be differentially methylated in alcohol abusers, but most evidence comes from studies that either had small sample size, were not specific to humans or were carried out using tissues other than blood **(22-33)**. Molecular mechanisms such as DNA methylation may underlie or enhance a predisposition to addictions and substance abuse, including alcohol drinking **(23, 27, 29, 31-34)**.

In the present study, we sought to (i) identify novel associations between alcohol consumption and blood DNA methylation, (ii) replicate previously reported associations, (iii) assess the reversibility of associations, and (iv) assess associations with changes in alcohol consumption using longitudinally collected data. We used samples from the Melbourne Collaborative Cohort Study (MCCS) to discover potential associations and sought to replicate the findings using samples from the Cooperative Health Research in the Augsburg Region (KORA) and London Life Sciences Prospective Population (LOLIPOP) studies.

## MATERIALS AND METHODS

### Study participants

Between 1990 and 1994 (baseline), 41,513 participants were recruited to the Melbourne Collaborative Cohort study (MCCS). The majority (99%) were aged 40 to 69 years and 41% were men. Southern European migrants were oversampled to extend the range of lifestyle-related exposures **(35).** Participants were contacted again between 2003 and 2007. Blood samples were taken at baseline and follow-up from 99% and 64% of participants, respectively. Baseline samples were stored as dried blood spots on Guthrie cards for the majority (73%), as mononuclear cell samples for 25% and as buffy coat samples for 2% of the participants. Follow-up samples were stored as buffy coat aliquots and dried blood spots on Guthrie cards.

All participants provided written informed consent and the study protocols were approved by the Cancer Council Victoria Human Research Ethics Committee.

The present study sample comprised MCCS participants selected for inclusion in one of seven previously conducted nested case-control studies of DNA methylation **(36-40)**. Controls were matched to incident cases of prostate, colorectal, gastric, lung or kidney cancer, urothelial cell carcinoma or mature B-cell neoplasms on: sex, year of birth, country of birth, baseline sample type and smoking status (the latter for the lung cancer study only). Participants included in each nested case-control study were free of cancer at baseline. After quality control, methylation data for baseline blood samples (baseline study) were available for 5,606 MCCS participants. Methylation measures were repeated in DNA extracted from blood samples collected on Guthrie cards at follow-up (longitudinal study) and, after quality control, were available for a subset of 1,088 of the controls who also had their baseline sample collected on a Guthrie card (***Table 1***).

**Table 1.** Characteristics of participants in the Melbourne Collaborative Cohort Study (MCCS) at baseline and follow-up visits.

|  | Cross-sectional analysis |  | Longitudinal analysis |  |  |  |
| --- | --- | --- | --- | --- | --- | --- |
|  | (all) |  | Baseline data |  | Follow-up data |  |
|  | (N=5,606) |  | (N=1,088) |  | (N=1,088) |  |
| Age in years, median [IQR] | 61 [54-65] |  | 59 [51-64] |  | 70 [63-76] |  |
| Sex, male | 3,793 | 68% | 740 | 68% | 740 | 68% |
| Country of birth |  |  |  |  |  |  |
| AU/NZ/Other | 3,744 | 67% | 831 | 76% | 831 | 76% |
| Greece | 434 | 8% | 44 | 4% | 44 | 4% |
| Italy | 819 | 15% | 89 | 8% | 89 | 8% |
| UK | 609 | 11% | 124 | 11% | 124 | 11% |
| BMI (kg/m²), median [range] | 26.9 [24.5-29.5] |  | 26.3 [24.1-29.0] |  | 26.8 [24.2-29.4] |  |
| Smoking status |  |  |  |  |  |  |
| Never | 2,519 | 45% | 549 | 50% | 545 | 50% |
| Former ≥15 years ago | 1,152 | 21% | 230 | 21% | 485 | 45% |
| Former <15 years ago | 1,100 | 20% | 196 | 18% |  |  |
| Current <20 cig/day | 322 | 6% | 62 | 6% | 58 | 5% |
| Current ≥20 cig/day | 513 | 9% | 51 | 5% |  |  |
| Drinking status |  |  |  |  |  |  |
| Lifetime abstainers | 1,314 | 24% | 209 | 20% | 168 | 16% |
| Former drinkers | 569 | 10% | 94 | 9% | 111 | 11% |
| Current drinkers | 3,626 | 66% | 759 | 71% | 744 | 73% |
| Median [IQR] for intake in last week (g/day) | 4.3 [0.0-18.7] |  | 4.3 [0.0-18.6] |  |  |  |
| Median [IQR] for lifetime intake (g/day) | 8.0 [0.3-23.0] |  | 8.1 [0.4-23.4] |  |  |  |
| Median [IQR] for intake in last year (g/day) |  |  |  |  | 7.9 [0.3-22.7] |  |
| Difference F-Up baseline (last week) (g/day) |  |  |  |  | 0.0 [-2.5 - 6.4] |  |
| Difference F-Up baseline (lifetime) (g/day) |  |  |  |  | 0.0 [-5.5 - 5.0] |  |
| Alcohol uptake (non-drinkers at baseline who were drinkers at follow-up) (yes) |  |  |  |  | 107 | 10% |
| Alcohol cessation (drinkers at baseline who were non-drinkers at follow-up) (yes) |  |  |  |  | 88 | 8% |

### Alcohol and other variables

At both baseline and follow-up, participants completed questionnaires that included detailed questions on demographic characteristics, medical history, cigarette smoking, alcohol consumption, physical activity and diet, the latter using food frequency questionnaires. On both occasions, anthropometric measurements were obtained by trained personnel using standard procedures. Height was only measured at baseline.

Alcohol intake at baseline was recorded as frequency and quantity of intake per drinking occasion by type (beer, wine, spirits) and by decade of age starting from 20 years. Participants were also asked about their alcohol intake on each day during the previous week, in terms of the number, measure and type of drink (e.g. two glasses of wine). At follow-up, the frequency and quantity of intake, by type of drink, during the previous calendar year were assessed as described above. Grams per day (g/day) were calculated as reported previously **(41)**. Participants who reported a weekly or current intake >200g/day were excluded. Four alcohol consumption variables were considered: g/day in the last week (continuous), g/day in the current decade (continuous), g/day over the lifetime (continuous), and drinking status (never, former, current; based on current decade and lifetime variables).

### DNA methylation and genetic data

Methods relating to DNA extraction and bisulfite conversion, DNA methylation data processing, normalisation and quality control, and genotyping are described in the ***Supplementary Methods***.

### Methylome-wide association study

We assessed cross-sectional associations at each individual CpG by regressing DNA methylation M-values on alcohol consumption using linear mixed-effects regression models, using the function *lmer* from the R package *lme4*. Alcohol intake was represented using three continuous variables (consumption in the previous year, consumption in the previous week, and lifetime consumption, in g/day) that were modelled separately. Models were adjusted by fitting fixed effects for age (continuous), sex, smoking status (never, former ≥15 years ago, former <15 years ago, current <20 cigarettes per day, current ≥20 cigarettes per day), BMI (≤25 kg/m^2^, >25 to ≤30, >30), country of birth (Australia/New-Zealand, Italy, Greece, United Kingdom), sample type (peripheral blood mononuclear cells, dried blood spots, buffy coats) and white blood cell composition (percentage of CD4+ T cells, CD8+ T cells, B cells, NK cells, monocytes and granulocytes, estimated using the Houseman algorithm **(42)**), and random effects for study, plate, and chip. A significance threshold of *P*-value<10^‒7^was used to account for multiple testing **(12)**. The False Discovery Rate (FDR) was used to identify suggestive associations.

Interaction terms were tested between alcohol consumption (previous week) and each of age (continuous), sex, smoking status (pseudo-continuous variable 1: Never, to 5: Current smoker >20 cigarettes/day), BMI (continuous), country of birth (as categorised above), future cancer case status, and a polygenic score for alcohol consumption, the latter derived as described below. These analyses were restricted to CpGs associated at P<10^‒7^ in the cross-sectional analysis with any of the three continuous alcohol variables (‘last week’, ‘current decade’ and ‘lifetime’ intake).

Associations for each type of alcohol (beer, wine, spirits) were assessed by including the three variables in a same model, so that intake of each type was adjusted for the two others. This was done for the ‘current decade’, and ‘lifetime’ alcohol intake variables because type of alcoholic beverage was not determined for the ‘alcohol last week’ variable.

Sensitivity analyses were conducted to assess potential confounding by: i) fitting the same models without adjustment for smoking or BMI; ii) fitting the same models with additional adjustment for socioeconomic status, educational attainment, a physical activity score based on metabolic equivalents **(43)**, and a score of healthy dietary habits **(44)**; and iii) examining whether methylation was associated with these covariates.

### Replication of novel associations

CpGs found to be associated with alcohol intake at P<10^‒7^ in the MCCS were selected for replication using data from the KORA cohort (N=1,662, assessed in 1999-2001) including German participants and the LOLIPOP cohort (N=4,042, assessed in 2003-2008), including predominantly Asian participants, respectively **(2, 45)**. Alcohol intake in KORA was defined as the average of ‘alcohol in previous day’ and ‘alcohol in previous week’. Alcohol intake in LOLIPOP was defined over a week (***Supplementary Methods***). Each cohort applied a normalisation method based on control probes **(46)** and adjusted models for the same covariates defined in a similar way to those used in the MCCS analyses, ***Supplementary Table 1, Supplementary Methods***. Results from the two cohorts were pooled using fixed-effects meta-analysis with inverse-variance weights **(47)**. An association was considered replicated if P<0.05 and the direction of association was the same as in the MCCS **(12)**.

### Polygenic score for alcohol consumption

A polygenic score for alcohol consumption was constructed using MCCS data based on the genome-wide association study by Clarke and colleagues, which identified 14 single nucleotide polymorphisms (SNPs) associated with alcohol consumption using UK Biobank data **(48)**. Data for 13 out of the 14 SNPs were available and were combined using the formula: *Polygenic score[i] = b_1_*d_1,i_ + … b_13_* d_13,i,_* where *d_k,i_* is the imputed allele dosage of variant *k* for person *i*, and *b_k_* the per-allele regression coefficient reported for SNP *k* **(59, 60)**.

### Replication of previously reported associations with alcohol consumption

Using MCCS data, we assessed replication (P<0.05) of associations with alcohol consumption (g/day) reported in a recent pooled, large-scale analysis of Europeans and African Americans **(21).** A total of 518 CpGs were considered, comprising 363 identified for participants of European ancestry participants and a further 155 CpGs identified for those of African ancestry.

### Reversibility of associations

We calculated regression coefficients for comparisons of ‘*former’* to ‘*never’*, ‘*current’* to ‘*never’*, and ‘*current’* to ‘*former’* drinkers using MCCS data. As there were too few never-drinkers in the KORA data (N=22), we only considered the comparison ‘*former’* to ‘*current’*; we pooled the latter using fixed-effects meta-analysis.

In the MCCS, we calculated a ‘reversibility coefficient’, expressed as a percentage defined as: coefficient (‘*former’* compared with ‘*current’*) / coefficient (‘*never’* compared with ‘*current’*). These analyses were undertaken for CpGs with *P*<10^‒7^ in the MCCS EWAS.

### Longitudinal associations

We further examined longitudinal associations with alcohol consumption for CpGs with *P*<10^‒7^ in the MCCS EWAS, incorporating data from follow-up. Linear mixed-effects regression models were used to assess the association between changes in DNA methylation (outcome) and changes in alcohol consumption (exposure) at each CpG. The change in alcohol consumption was computed in g/day as the difference between follow-up (alcohol intake in previous year) and baseline (previous week intake); study was included as a random effect and the following variables were included as fixed effects: baseline alcohol intake, baseline BMI and change in BMI (continuous), baseline age and change in age (continuous), sex, smoking status at baseline (as defined previously), smoking status at follow-up (yes/no), country of birth (as defined previously), baseline cell composition (as defined previously), change in each cell type composition (continuous) and baseline methylation M-value. The change in methylation was calculated as the difference between follow-up and baseline ComBat-normalised methylation M-values. The same analyses were conducted for the KORA cohort, in which methylation measures taken approximately seven years later (2006-2008) were available for 1,332 participants (***Supplementary Methods***). As adjustment for baseline methylation in analyses of change in methylation may lead to bias in some circumstances **(49)**, we conducted a sensitivity analysis using models without adjustment for baseline methylation in the MCCS.

### Pathway analyses

We used the *gometh* function from the *missMethyl* package **(50)** for pathway analyses assessing over-representation relative to all KEGG pathways **(51)**. To investigate potentially different biological pathways underlying i) acute compared with chronic alcohol-associated consumption and ii) most dynamic compared with least dynamic methylation sites, *gometh* was applied, respectively, to i) CpGs associated with ‘last week’ and ‘lifetime’ alcohol intake, and ii) CpGs with a reversibility coefficient greater and lower than 50%. A *P*-value lower than 0.05 was considered to indicate a potentially relevant pathway.

All statistical analyses were performed using the software R (version 3.4.0).

## RESULTS

Altogether, 5,606 MCCS participants were included in the cross-sectional analysis; their median age was 61 years (IQR: 54-65), 68% were males, and alcohol intakes were wide-ranging (***Table 1***). There were moderate-to-high correlations between the baseline alcohol variables, as well as between the baseline and follow-up variables, with Spearman correlations ranging from 0.68 to 0.76 (***Supplementary Table 2***). Participants in the longitudinal analysis were younger and generally had healthier lifestyle.

### Methylome-wide association study

Across the three cross-sectional continuous alcohol variables considered, we observed 1,414 associations with *P*<10^‒7^. The most statistically significant associations are presented in ***Table 2***. There were 1,318, 358, and 392 CpGs associated with alcohol intake over the last week, decade and lifetime, respectively ***(Supplementary Table 3, Supplementary Figure 1)***. Associations were consistently stronger for alcohol intake in the previous week (***Supplementary Figure 2***).

**Table 2.**
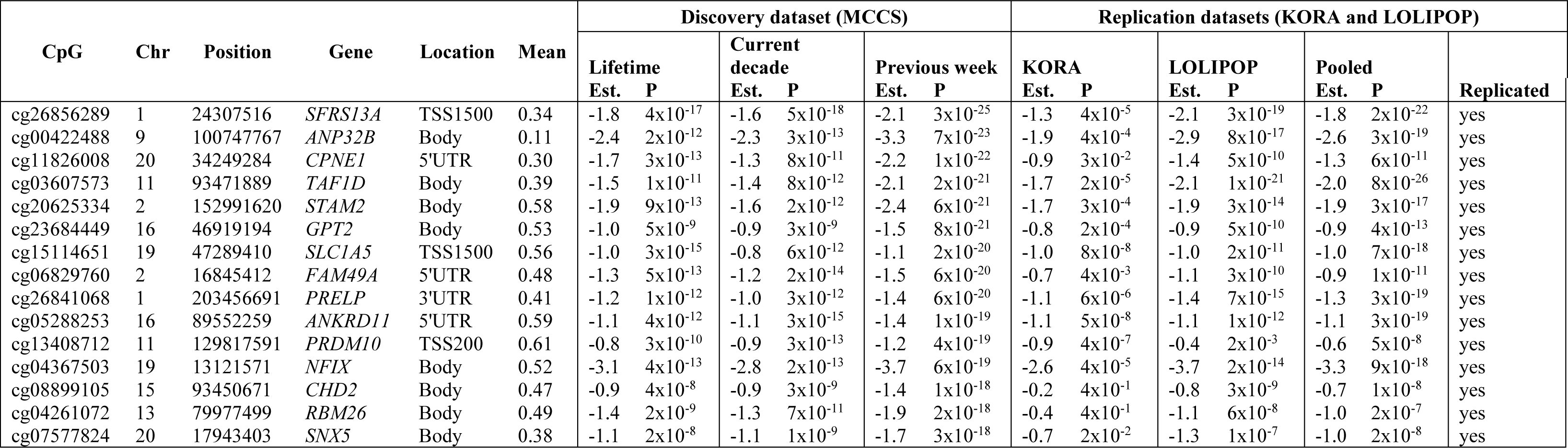
The 15 most statistically significant novel associations (N=1,243) between alcohol intake and blood DNA methylation discovered in the MCCS and replication in external cohorts KORA and LOLIPOP.

<u>Abbreviations</u>: Chr.: Chromosome; Est.: regression coefficients from linear mixed regression models (MCCS) or linear regression models (KORA and LOLIPOP).

All models were adjusted by fitting fixed effects for age, sex, smoking status, BMI, country of birth, sample type and white blood cell composition (percentage of CD4+ T cells, CD8+ T cells, B cells, NK cells, monocytes and granulocytes, estimated using the Houseman algorithm), and batch effects.

Est. pooled and P pooled are results from the fixed-effects meta-analysis of results from KORA and LOLIPOP.

Regression coefficients are given for intakes in grams per day and multiplied by 1000.

The full results are presented in the Supplementary file.

Of these 1,414 associations, 1,243 were novel (i.e. not reported in Liu et al. **(21)**), and the CpGs were located in 831 genes; 241 CpGs were intergenic. Compared to the rest of the HM450 assay, alcohol-associated CpGs were over-represented in gene bodies (binomial proportion test, *P*=0.001), promoter regions TSS1500 (*P*=5×10^‒6^) and 5’UTR (*P*=3×10^‒8^); unannotated regions were underrepresented (*P*=2×10^‒5^). These 1,414 associations corresponded to 1,084 unique clusters when considering that CpGs separated by less than 50kb of each other formed a single genomic region (***Supplementary Table 4***). For the overwhelming majority (99%) of associations, greater alcohol intake was associated with lower methylation. The number of CpGs associated with alcohol intake over the last week, decade and lifetime was 16,732, 5,751 and 6,585, respectively when considering an FDR-adjusted *P*<0.05 (***Supplementary Table 3***).

#### Replication of novel associations using external data

Of the 1,243 novel associations, 1,078 (87%) were replicated using data from KORA and LOLIPOP. Replication rates were 87%, 89% and 93% for CpGs associated with alcohol intake over the last week, decade and lifetime, respectively (P<0.05; ***Table 2*** and ***Supplementary Table 5***).

#### Interaction analyses

Using the Bonferroni correction for multiple testing (P=0.05/1,414=3.5×10^‒5^) and the ‘last week’ alcohol intake variable, we observed stronger associations for women at cg13446906 (*MIR548F5*), and cg22363327 (*SFRS13B*), and weaker associations for smokers at cg05104080 (*ILKAP*), cg17058475 (*CPT1A*), and cg01395047 (*TLR9*). At P<0.05, stronger associations were observed for women at 200 CpGs (test for binomial proportions, P=6×10^‒29^); weaker associations were observed for participants with a higher smoking score (N= 159, P=6×10^‒20^) and a higher BMI (N=165, P=0.003) (***Table 3***).

**Table 3.**
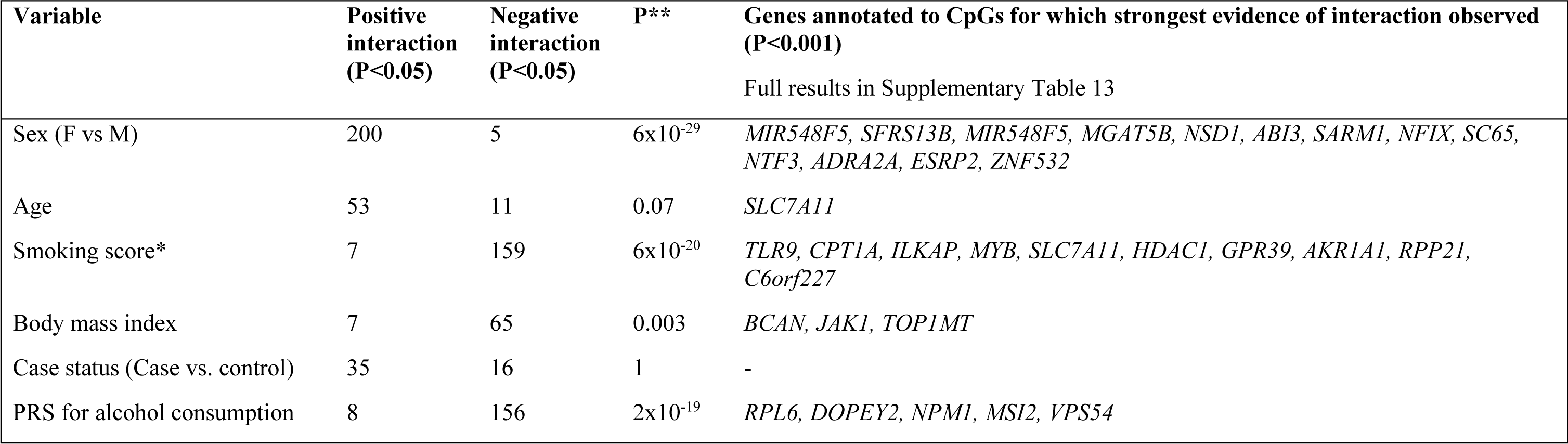
Interaction between alcohol intake and other factors in association with DNA methylation changes for 1,414 alcohol-related CpG sites.

a Smoking was coded as a pseudo-continuous score: 0: Never, 1: Former, ≥15 years ago, 2: Former <15 years ago, 3: Current <20 cig/day, 4: Current ≥20 cig/day.

** *P*-value from test of binomial proportions that the greatest number (e.g. 200 for sex) is greater than 35 (number of associations expected by chance), assuming independence between CpGs.

For the MCCS sample with genetic and DNA methylation data (N=3,859), the polygenic score was positively associated with alcohol intake in previous week (P=9×10^‒5^) (***Supplementary Table 12***). Weaker associations (P<0.05) for participants with higher polygenic score were observed for 156 CpGs (P=2×10^‒19^), although none passed the Bonferroni correction (***Table 3 and Supplementary Tables 12-13***).

#### Alcohol types

For alcohol intake in the previous decade, the regression coefficients were the greatest for beer intake for 917 (65%) CpGs, for spirit intake for 309 (22%) CpGs and for wine intake for 188 (13%) CpGs, whereas these percentages were 27%, 32% and 41%, respectively, for lifetime alcohol intake (***Supplementary Table 14***).

#### Sensitivity analyses

Age, sex, smoking, BMI, country of birth, and cell composition were associated with methylation at many alcohol-associated CpG sites. Adjustment for smoking, but not BMI, made a substantial difference to the estimated coefficients; for the ‘previous week’ alcohol intake variable, 1,985 associations were observed when no adjustment for smoking was made. Adjustment for additional health-related variables made virtually no difference to the results (***Supplementary Material***).

### Replication of previously reported associations using MCCS data

We examined the replication of the associations between alcohol intake and whole-blood DNA methylation previously reported in Liu et al. with P<10^‒7^ **(21).** Of the 518 associations, we replicated 403 (78%) at P<0.05, using the MCCS ‘previous week’ alcohol intake variable; 169 reached genome-wide significance P<10^‒7^ (***Table 4, Supplementary Table 5***). Replication was substantially higher for associations identified in European-ancestry individuals (335/363, 92%), compared with those of African ancestry (68/155, 44%).

**Table 4.**
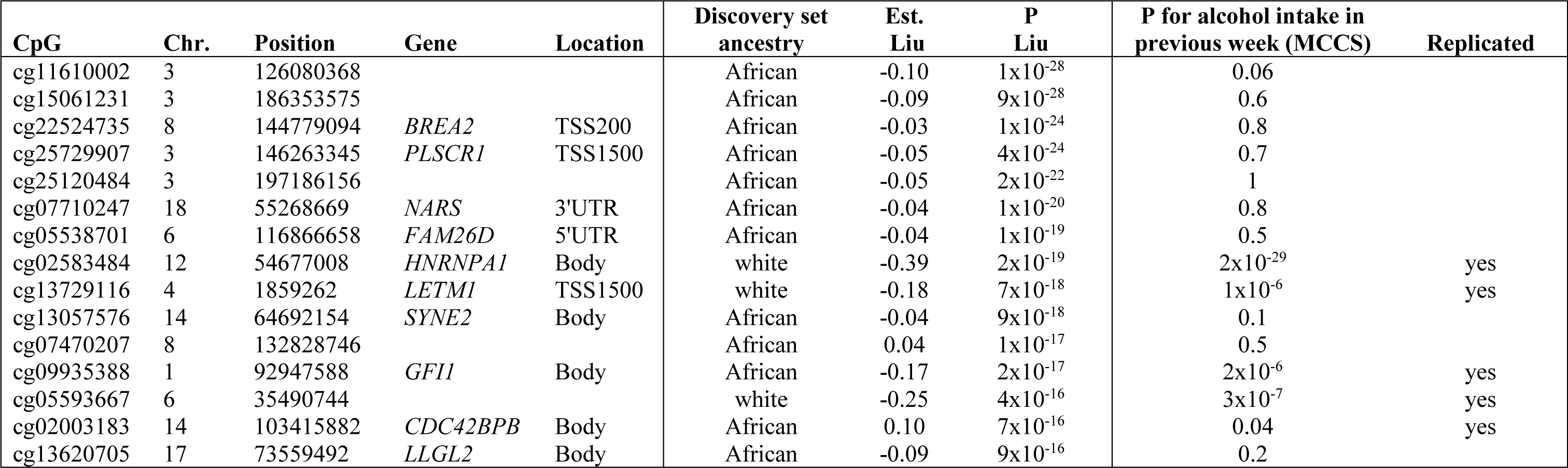
Replication of the strongest associations (P<10^‒15^) reported in Liu et al., Mol. Psychiatry, 2016 using MCCS data.

Abbreviations: Chr.: chromosome; Est. Liu: regression coefficient reported in the study by Liu et al., Mol. Psychiatry, 2016

The full results are presented in the Supplementary file.

### Reversibility of associations

The ‘current decade’ alcohol consumption variable of the MCCS was classified into current, former, and never. Of the 1,414 CpGs considered, 280 were differentially methylated (*P*<0.05) between former and never drinkers, and 282 were differentially methylated (*P*<0.05) between former and current drinkers, with only 2 overlapping CpGs. The reversibility coefficients comparing former to never and current to never drinkers were wide-ranging (median: 47%, IQR=19% to 79%) (***Supplementary Table 7***). In KORA, there was also substantial reversibility, as 332 CpGs were differentially methylated when comparing former to current drinkers. A total of 86 CpGs were differentially methylated in both datasets for the comparison of former to current drinkers, and 530 CpGs when pooling results from both cohorts (P<0.05); the most significant associations are presented in ***Table 5***.

**Table 5.** Reversibility of associations (cross-sectional Current vs Former vs Never) for the fifteen most significant associations in the MCCS EWAS for the ‘current decade’ alcohol intake variable.

| CpG | Chr. | Position | Current decade intake (g/day) |  | Former vs. never |  | Current vs. never |  | Reversibility coefficient |
| --- | --- | --- | --- | --- | --- | --- | --- | --- | --- |
|  |  |  | Est. | P | Est. | P | Est. | P |  |
| cg06690548 | 4 | 139162808 | -6.6 | $7 \times 10^{-70}$ | -0.01 | 0.8 | -0.12 | $2 \times 10^{-8}$ | 95% |
| cg14476101 | 1 | 120255992 | -3.4 | $6 \times 10^{-34}$ | -0.02 | 0.4 | -0.10 | $2 \times 10^{-10}$ | 81% |
| cg12825509 | 3 | 185648568 | -2.4 | $2 \times 10^{-26}$ | -0.05 | 0.009 | -0.07 | $2 \times 10^{-8}$ | 31% |
| cg02711608 | 19 | 47287964 | -2.1 | $2 \times 10^{-24}$ | -0.01 | 0.5 | -0.04 | $8 \times 10^{-5}$ | 76% |
| cg18120259 | 6 | 43894639 | -1.9 | $1 \times 10^{-20}$ | -0.04 | 0.009 | -0.05 | $6 \times 10^{-6}$ | 14% |
| cg18336453 | 6 | 43082296 | -1.3 | $2 \times 10^{-19}$ | -0.03 | 0.01 | -0.04 | $5 \times 10^{-8}$ | 32% |
| cg19693031 | 1 | 145441552 | -2.5 | $5 \times 10^{-19}$ | -0.02 | 0.3 | -0.06 | $3 \times 10^{-4}$ | 61% |
| cg11376147 | 11 | 57261198 | -1.5 | $1 \times 10^{-18}$ | -0.04 | 0.001 | -0.05 | $3 \times 10^{-7}$ | 7% |
| cg26856289 | 1 | 24307516 | -1.6 | $5 \times 10^{-18}$ | -0.01 | 0.5 | -0.05 | $4 \times 10^{-6}$ | 78% |
| cg17058475 | 11 | 68607737 | -3.6 | $9 \times 10^{-18}$ | -0.07 | 0.03 | -0.16 | $5 \times 10^{-13}$ | 56% |
| cg16246545 | 1 | 120255941 | -1.7 | $1 \times 10^{-17}$ | 0.00 | 0.9 | -0.04 | $1 \times 10^{-4}$ | 104% |
| cg06644515 | 1 | 173834831 | -1.6 | $6 \times 10^{-17}$ | 0.00 | 0.9 | -0.04 | $3 \times 10^{-4}$ | 106% |
| cg02583484 | 12 | 54677008 | -1.4 | $1 \times 10^{-16}$ | -0.02 | 0.2 | -0.05 | $3 \times 10^{-7}$ | 62% |
| cg15804598 | 17 | 43224418 | -1.1 | $1 \times 10^{-15}$ | -0.02 | 0.1 | -0.04 | $2 \times 10^{-7}$ | 52% |
| cg00252472 | 6 | 150739173 | -2.7 | $1 \times 10^{-15}$ | -0.03 | 0.3 | -0.08 | $5 \times 10^{-6}$ | 63% |

Abbreviations: Chr.: Chromosome; Est.: regression coefficients from linear mixed regression models (MCCS) or linear regression models (KORA and LOLIPOP).

All models were adjusted by fitting fixed effects for age, sex, smoking status, BMI, country of birth, sample type and white blood cell composition (percentage of CD4+ T cells, CD8+ T cells, B cells, NK cells, monocytes and granulocytes, estimated using the Houseman algorithm), and batch effects.

Est. pooled and P pooled are results from the fixed-effects meta-analysis of results from KORA and LOLIPOP. Regression coefficients are given for intakes in grams per day and multiplied by 1000.

The full results are presented in the Supplementary file.

### Longitudinal associations

We further tested the 1,414 CpGs with cross-sectional associations for longitudinal associations using the MCCS and KORA. Repeated methylation measures and alcohol information were available a median of 11 and 7 years apart in the MCCS and KORA, respectively. Change in alcohol intake was associated with change in methylation (P<0.05) for 267 CpGs in the MCCS, and for 331 CpGs in KORA, with 92 overlapping associations. After pooling the results, we observed evidence of change over time for 513 CpG sites (***Supplementary Table 8***). The most statistically significant longitudinal associations are shown in ***Table 6***. Fewer associations were observed in the MCCS when no adjustment for baseline methylation levels was made (N=125, ***Supplementary Material***). The analyses of alcohol cessation (N=88 participants, 8%) and uptake (N=107, 10%) revealed 147 and 40 associations, respectively, overlapping little with the 513 identified CpGs (N=65 and N=19, respectively). CpG sites that showed stronger evidence of association in the longitudinal analysis appeared somewhat more reversible in the cross-sectional analysis (***Supplementary Figure*** 5) and 245 CpGs appeared differentially methylated in both analyses (***Supplementary Table 10***).

**Table 6.** Longitudinal associations (P<10^‒5^) assessed in the MCCS and in KORA (data collected 11 and 7 years apart, respectively).

| CpG | Chr. | Position | Gene | Location | Cross-sectional data |  | Longitudinal data |  |  |  |  |  |
| --- | --- | --- | --- | --- | --- | --- | --- | --- | --- | --- | --- | --- |
|  |  |  |  |  | MCCS (previous week) |  | MCCS |  | KORA |  | Pooled |  |
|  |  |  |  |  | Est. | P | Est. | P | Est. | P | Est. | P |
| cg06690548 | 4 | 139162808 | *SLC7A11* | Body | -7.9 | $1 \times 10^{-86}$ | -4.7 | $2 \times 10^{-5}$ | -5.4 | $7 \times 10^{-11}$ | -5.2 | $5 \times 10^{-15}$ |
| cg18120259 | 6 | 43894639 | *LOC100132354* | Body | -2.4 | $4 \times 10^{-27}$ | -1.5 | $5 \times 10^{-3}$ | -1.5 | $5 \times 10^{-8}$ | -1.5 | $8 \times 10^{-10}$ |
| cg02711608 | 19 | 47287964 | *SLC1A5* | 1stExon | -2.6 | $6 \times 10^{-30}$ | -2.0 | $4 \times 10^{-4}$ | -1.4 | $1 \times 10^{-6}$ | -1.5 | $3 \times 10^{-9}$ |
| cg14476101 | 1 | 120255992 | *PHGDH* | Body | -4.0 | $2 \times 10^{-39}$ | -2.9 | $1 \times 10^{-4}$ | -1.8 | $3 \times 10^{-6}$ | -2.1 | $3 \times 10^{-9}$ |
| cg16246545 | 1 | 120255941 | *PHGDH* | Body | -2.2 | $4 \times 10^{-23}$ | -1.7 | $2 \times 10^{-3}$ | -1.4 | $7 \times 10^{-7}$ | -1.4 | $4 \times 10^{-9}$ |
| cg11376147 | 11 | 57261198 | *SLC43A1* | Body | -2.1 | $2 \times 10^{-29}$ | -1.0 | $8 \times 10^{-3}$ | -1.4 | $1 \times 10^{-6}$ | -1.3 | $3 \times 10^{-8}$ |
| cg20732160 | 3 | 48590040 | *PFKFB4* | Body | -1.7 | $3 \times 10^{-16}$ | -1.6 | $3 \times 10^{-3}$ | -1.2 | $1 \times 10^{-5}$ | -1.3 | $1 \times 10^{-7}$ |
| cg03068497 | 7 | 30635838 | *GARS* | Body | -3.0 | $3 \times 10^{-14}$ | -2.0 | $2 \times 10^{-2}$ | -2.4 | $4 \times 10^{-6}$ | -2.3 | $2 \times 10^{-7}$ |
| cg07626482 | 19 | 47289503 | *SLC1A5* | TSS1500 | -1.4 | $4 \times 10^{-20}$ | -1.2 | $2 \times 10^{-3}$ | -1.0 | $4 \times 10^{-5}$ | -1.1 | $3 \times 10^{-7}$ |
| cg13526915 | 14 | 24164078 | | | -2.1 | $8 \times 10^{-13}$ | -1.2 | $8 \times 10^{-2}$ | -1.7 | $1 \times 10^{-5}$ | -1.6 | $3 \times 10^{-6}$ |
| cg14756878 | 2 | 12568736 | | | -1.2 | $3 \times 10^{-9}$ | -1.4 | $2 \times 10^{-3}$ | -0.9 | $4 \times 10^{-4}$ | -1.0 | $3 \times 10^{-6}$ |
| cg04460609 | 4 | 16532808 | *LDB2* | Body | -2.2 | $3 \times 10^{-19}$ | -1.2 | $2 \times 10^{-2}$ | -1.3 | $1 \times 10^{-4}$ | -1.3 | $5 \times 10^{-6}$ |
| cg21626848 | 17 | 39969267 | *SC65* | TSS1500 | -1.6 | $1 \times 10^{-16}$ | -1.8 | $2 \times 10^{-4}$ | -0.8 | $2 \times 10^{-3}$ | -1.1 | $8 \times 10^{-6}$ |
| cg03533472 | 16 | 46919112 | *GPT2* | Body | -2.2 | $1 \times 10^{-15}$ | -2.5 | $9 \times 10^{-4}$ | -1.7 | $2 \times 10^{-3}$ | -2.0 | $9 \times 10^{-6}$ |

Abbreviations: Chr.: Chromosome; Est.: regression coefficients from linear mixed regression models (MCCS) or linear regression models (KORA and LOLIPOP).

All models were adjusted by fitting fixed effects for age, sex, smoking status, BMI, country of birth, sample type and white blood cell composition (percentage of CD4+ T cells, CD8+ T cells, B cells, NK cells, monocytes and granulocytes, estimated using the Houseman algorithm), and batch effects.

Est. pooled and P pooled are results from the fixed-effects meta-analysis of results from KORA and LOLIPOP. Regression coefficients are given for intakes in grams per day and multiplied by 1000.

The full results are presented in the Supplementary file.

### Pathway analyses

The *gometh* function was applied separately to CpG sites associated with ‘lifetime’ (N=392) and ‘last week’ alcohol intake (N=1,318), and for associations with reversibility coefficients lower and greater than 50% (N=753 and N=661, respectively) (***Supplementary Table 11***). The most significant KEGG pathways were for the ‘last week’ variable: ‘*Chronic myeloid leukemia*’, ‘*Ribosome*’, ‘*Glycosaminoglycan biosynthesis*’; lifetime variable: ‘*Regulation of actin cytoskeleton*’, ‘*Biosynthesis of amino acids*’, ‘*Platelet activation*’; persistent associations: ‘*Ribosome’*, ‘*Human cytomegalovirus infection*’; reversible associations: “*Cellular senescence*”, “*MAPK signaling pathway*”. A few nominally significant associations were observed for other pathways directly relevant to alcohol drinking such as ‘*GABAergic synapse*’.

## DISCUSSION

Our study identified 1,481 methylation sites associated with alcohol consumption, including 1,078 discovered in the MCCS and replicated in independent cohorts and 403 replicated from a previous large EWAS. An additional 513 CpGs discovered in the cross-sectional analysis were indirectly replicated in the longitudinal analysis. The findings using a less conservative significance threshold (FDR) indicate that many more alcohol-associated CpGs are likely to exist across the genome. The majority of CpG sites we identified were hypomethylated with increased alcohol intake.

Alcohol-related hypomethylation appears to be largely reversible upon alcohol cessation; this was inferred from three analyses. First, we observed substantially more and stronger associations with alcohol consumed in the last week than in the last decade or lifetime, indicating that the alcohol intake most relevant to DNA methylation was that closest to blood draw. Similar data were not available from other cohorts to replicate this finding. Second, the cross-sectional comparison of current and former to never drinkers revealed that the difference in terms of DNA methylation between former and current drinkers was on average half that between never and current drinkers, with wide-ranging estimates; we also identified CpG sites that were consistently differentially methylated in former compared with current drinkers in MCCS and KORA. Third, using longitudinal data taken several years apart (11 years in the MCCS and 7 years in KORA), we identified a set of 513 CpG sites that varied with change in alcohol consumption, and 245 of these corresponded to differentially methylated sites in the comparison of former to current drinkers (cross-sectional). The longitudinal analysis had less power due to a lower number of included participants and relatively small variation in drinking status over the periods considered, and because the variable reflecting changes in alcohol drinking may have been measured with error. These findings taken together indicate a substantial degree of reversibility in the associations, which was not assessed by previous studies.

Another potential limitation of our study is residual confounding, most notably by smoking or white blood cell type composition, which are both strongly associated with alcohol drinking and DNA methylation **(15)**. We observed that many CpG sites associated with alcohol drinking were also associated with other factors such as smoking, white blood cell composition, BMI and other factors, which may indicate that these loci are very sensitive to the environment. Cell composition was estimated with the widely used Houseman algorithm modified by Jaffe and Irizarry **(42, 52)** and we did not assess sensitivity to the method used for deriving cell composition **(53)**. Although our adjustment for smoking was relatively comprehensive, our sensitivity analyses demonstrate that alcohol and smoking may exert joint influences on many CpGs across the genome.

We observed less substantial replication for CpGs discovered in individuals of African ancestry, which may indicate that alcohol-associated methylation changes are not generalizable to all human populations. Associations in individuals of African ancestry in the study by Liu et al. were discovered by pooling data from a smaller number of studies so might have been less replicable by nature.

The associations between alcohol consumption and methylation were stronger for people with genetic predisposition to consume less alcohol, as defined by a 13-SNP polygenic score. Several genes included in the polygenic score are involved in alcohol metabolism, for example those of the alcohol dehydrogenase family (*ADH1B, ADH1C,* and *ADH5)* and *GCKR* (glucokinase regulatory protein, involved in glucose metabolism), which could provide a biological explanation for the interaction between the polygenic score and alcohol intake, given links between alcohol metabolism pathway and epigenetic mechanisms **(54)**. The polygenic score explained 0.5% of variance in total alcohol consumption, which is consistent with other studies (0.6% in **(48),** and 0.11% in **(55)**). In comparison, the predictors of alcohol consumption presented in Liu et al. explained 5-10% and 12-14% of variance with 5 and 144 CpGs, respectively **(21).**

Associations appeared weaker for smokers and men, consistent with the observation that these population subgroups tend to drink more alcohol in most cultures, perhaps due to being less susceptible to the harmful effects of alcohol **(56).** Women have previously been reported to have slower alcohol metabolism than men **(57)**. These findings should be confirmed by further studies. We included in the analysis participants who later developed cancer, which could give rise to collider bias when both DNA methylation and alcohol are associated with cancer risk **(58)**. We found no evidence of differences in associations by case/control status in our study. Further, that most discovered associations were replicated in independent cohorts of healthy participants with distinct ethnic origin is a strong testament that our findings were not driven by the inclusion of future cancer cases.

The newly discovered CpG sites with strongest evidence of association with alcohol consumption were all located in genes, including in the regulatory regions of, for example, *SFRS13A, CPNE1*, *SLC1A5, FAM49A, PRELP, ANKRD11, PRDM10, COMT*, on which to our knowledge little research has been conducted in relation to alcohol metabolism or consumption. Some studies of alcohol-associated methylation changes have used tissues other than blood, particularly from the brain, and reported that DNA methylation might be the cause rather than the consequence of alcohol consumption, at least at certain loci. We did not examine causality in our study; we hypothesise that if DNA methylation were the cause of alcohol drinking, it would likely be at a restricted number of loci involved in addiction mechanisms and alcohol metabolism. We did not identify strongly enriched biological pathways that are key to alcohol metabolism or alcohol-related diseases such as cancer, cardiovascular disease, and mental health and addiction pathologies.

Our study shows that alcohol consumption is associated with widespread changes in blood DNA methylation. These changes appear more pronounced in women, non-smokers, and individuals with lower genetic predisposition to drink alcohol, and are at least partially reversible.

## ACKNOWLEDGEMENTS

<u>MCCS:</u> This work was supported by the Australian National Health and Medical Research Council (NHMRC) [grant 1088405]. MCCS cohort recruitment was funded by VicHealth and Cancer Council Victoria. The MCCS was further supported by Australian NHMRC grants 209057, 251553 and 504711 and by infrastructure provided by Cancer Council Victoria. Cases were ascertained through the Victorian Cancer Registry (VCR) and the Australian Cancer Database (Australian Institute of Health and Welfare). The nested case-control methylation studies were supported by the NHMRC grants 1011618, 1026892, 1027505, 1050198, 1043616 and 1074383. M.C.S. is an NHMRC Senior Research Fellow (1061177).

<u>KORA</u>: The KORA study was initiated and financed by the Helmholtz Zentrum München – German Research Center for Environmental Health, which is funded by the German Federal Ministry of Education and Research (BMBF) and by the State of Bavaria. Furthermore, KORA research has been supported within the Munich Center of Health Sciences (MC-Health), Ludwig-Maximilians-Universität, as part of LMUinnovativ. This work has received funding from the European Foundation for Alcohol Research (ERAB 2018 – EA1817). We thank all members of field staffs who were involved in the planning and conduct of the MONICA/KORA Augsburg studies.

<u>LOLIPOP</u>: The LOLIPOP study is supported by the National Institute for Health Research (NIHR) Comprehensive Biomedical Research Centre Imperial College Healthcare NHS Trust, the British Heart Foundation (SP/04/002), the Medical Research Council (G0601966, G0700931), the Wellcome Trust (084723/Z/08/Z, 090532 & 098381) the NIHR (RP-PG-0407-10371), the NIHR Official Development Assistance (ODA, award 16/136/68), the European Union FP7 (EpiMigrant, 279143) and H2020 programs (iHealth-T2D, 643774). We acknowledge support of the MRC-PHE Centre for Environment and Health, and the NIHR Health Protection Research Unit on Health Impact of Environmental Hazards. The work was carried out in part at the NIHR/Wellcome Trust Imperial Clinical Research Facility. The views expressed are those of the author(s) and not necessarily those of the Imperial College Healthcare NHS Trust, the NHS, the NIHR or the Department of Health. We thank the participants and research staff who made the study possible. JC is supported by the Singapore Ministry of Health’s National Medical Research Council under its Singapore Translational Research Investigator (STaR) Award (NMRC/STaR/0028/2017).

## AUTHORS CONTRIBUTION

PAD, JEJ, EM, DRE, GGG, and RLM were responsible for the study concept and design. CHJ, JEJ, EM, DFS, LB, GS, CG, KHL, AP, JSK, MCS, DRE, MW, JCC, GGG, and RLM contributed to the acquisition of data. HJ, XW, CHJ, and JEJ contributed to the preparation of the data. PAD, RW, and BL performed the statistical analyses. PAD and RLM drafted the manuscript. All authors provided critical revision of the manuscript for important intellectual content. All authors critically reviewed content and approved final version for publication.

